# Distinguish between Duplication of Essential Genes and Duplication of Dispensable Genes

**DOI:** 10.1101/2020.03.10.986257

**Authors:** Xun Gu

## Abstract

When a dispensable gene is duplicated (ancestral dispensability), genetic buffering and duplicate compensation together maintain the gene dispensability, whereas duplicate compensation is the only mechanism when an essential gene is duplicated (ancestral essentiality). To explore the distinct pattern of genetic robustness between these evolutionary scenarios, we formulated a probabilistic model with some biologically reasonable assumptions for analyzing a set of duplicate pairs with three possible states: double-dispensable (DD), semi-dispensable (one dispensable one essential, DE) or double-essential (EE). A computational pipeline is then developed to predict the distribution of three states (DD, DE and EE) conditional of ancestral dispensability or essentiality, respectively. This model was applied to yeast duplicate pairs from a whole-genome duplication, revealing that the process of essentiality of those duplicated from essential genes could be significantly higher than that of those duplicated from dispensable genes. We thus proposed a hypothesis that the process of sub-functionalization may be faster than neo-functionalization. Our analysis may provide some new insights about the role of duplicate compensation on genetic robustness.

## Introduction

The role of functional compensation by duplicate genes has been examined in diverse organisms by comparing the proportion (*P_E_*) of essential genes in duplicates to *P_E_* in singletons (Conant and Wagner 2004; Gu, et al. 2003; Hanada, et al. 2009; Liang and Li 2007; Liao and Zhang 2007; Wagner 2000). If duplicates play a significant role in functional compensation, the *P_E_* for duplicates should be lower than that of singletons. While this is indeed the case in yeasts, worms, and plants (Conant and Wagner 2004; Gu, et al. 2003; Hanada, et al. 2009; Hanada, et al. 2011; Qian, et al. 2010), no significant difference in *P_E_* was found between mouse single-copy and duplicate genes (Liang and Li 2007; Liao and Zhang 2007). A number of explanations were proposed. For instance, Su and Gu (2008) noticed that most mouse knockout duplicates tend to be ancient, subject to the loss of duplicate compensation by the mechanism of subfunctionalization. Meanwhile, some studies showed that functional and protein connectivity bias between essential and dispensable duplicate genes may be the cause (Makino, et al. 2009; Liang and Li 2009).

The pattern of duplicate compensation is complex (Diss, et al. 2017; Hahn 2009; Keane, et al. 2014; Szklarczyk, et al. 2008; Teufel, et al. 2018). When an essential gene is duplicated (termed *ancestral essentiality*), duplicate compensation is the only mechanism to keep two resulting copies dispensable. On the other hand, when a dispensable gene is duplicated (termed *ancestral dispensability*), the ancient genetic buffering and duplicate compensation together keep both duplicate copies dispensable. However, all *P_E_*-related analyses in the literature did not distinguish between these two possibilities.

This paper will address this issue as follows. We first develop a statistical method to analyze a set of duplicate pairs with three possible states: double-dispensable (DD), semi-dispensable (one dispensable one essential, DE) or double-essential (EE). With the help of some biologically reasonable assumptions, a probabilistic model is formulated, which takes the duplication of an essential gene or a dispensable gene into account. A computational pipeline is then developed to predict the distribution of three states (DD, DE and EE) conditional of ancestral dispensability or essentiality, respectively. Exemplified by the yeast duplicate pairs from the whole genome duplication (WGD), the new model compared the proportion of essential genes between those duplicated from essential genes and those dispensable genes, and discussed new insights for the evolutionary pattern of genetic robustness after gene duplication.

## Results and Discussion

### Genetic robustness between duplicate genes

A gene is called ‘*essential*’ (denoted by *d*^-^) if the single-gene deletion phenotype is severe or lethal, or ‘*dispensable*’ (denoted by *d*^+^) if its deletion phenotype is normal or nearly-normal (Hsiao and Vitkup 2008; Ihmels, et al. 2007; Su and Gu 2008). Consider two paralogous genes (*A* and *B*) duplicated from a common ancestor (*O*) *t* time units ago. There are four combined states, denoted by (*d_A_, d_B_*), representing double-dispensable (*d*^+^,*d*^+^), semi-dispensable (*d*^+^,*d*^-^) or (*d*^-^,*d*^+^), or double-essential (*d*^-^,*d*^-^), respectively.

First consider *P_t_*(*d_A_, d_B_*), the probability of any joint states (*d_A_, d_B_*) at time *t*. To derive *P_t_*(*d_A_, d_B_*), one should distinguish between the duplication of an essential genes (ancestral essentiality, denoted by *O*^-^) and the duplication of a dispensable gene (ancestral dispensability, denoted by *O*^+^). This idea can be statistically formulated as follows. Let *Q_t_*(*d_A_, d_B_*|*O*^-^) be the probability of being (*d_A_, d_B_*) after *t* time units since gene duplication, conditional of the ancestral essentiality (*O*^-^), and *Q_t_*(*d_A_*, *d_B_*|*O*^+^) be the probability conditional of the ancestral dispensability (*O*^+^). Since the ancestral state (dispensable or essential) for a duplicate pair is usually unknown, a mixture model is implemented as follows. Let *R_0_*=*P*(*O*^+^) be the probability of a gene pair duplicated from a dispensable gene, and 1-*R_0_*=*P*(*O*^-^) be that from an essential gene. Together, one can write

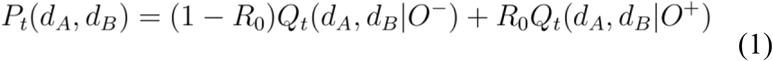

where (*d_A_*, *d_B_*)= (*d*^+^, *d*^+^), (*d*^+^, *d*^-^), (*d*^-^, *d*^+^) or (*d*^-^, *d*^-^), respectively.

Suppose we have a set (*N*) of duplicate pairs; all *2N* genes have single-gene deletion phenotypes (dispensable or essential). Three types of duplicate pairs are considered, that is, *DD* for (*d*^+^,*d*^+^), *DE* for (*d*^+^, *d*^-^) or (*d*^-^, *d*^+^), and *EE* for (*d*^-^, *d*^-^), and their frequencies are denoted by *f_DD_, f_DE_* and *f_EE_*, respectively. An intriguing question we wish to address is how we can estimate *Q_t_*(*d_A_, d_B_*|*O*^-^) and *Q_t_*(*d_A_, d_B_*|*O*^+^) from observed frequencies *f_DD_, f_DE_* and *f_EE_*. As shown below, to this end we develop a three-step approach: first develop simple models for; second estimate or specify model parameters from the observations; and third calculate *Q_t_*(*d_A_, d_B_*|*O*^-^) and *Q_t_*(*d_A_, d_B_*|*O*^+^).

### Sub-functionalization model of *Q_t_*(*d_A_*, *d_B_*|*O*^-^)

When an essential gene was duplicated, the process of *sub-functionalization* has been thought to be the major evolutionary mechanism for duplicate preservation (Innan and Kondrashov 2010; Prince and Pickett 2002; Stark, et al. 2017). As a result, both duplicate copies can be preserved without the need to invoke positive selection. Here we develop a simple model for *Q_t_*(*d_A_, d_B_*|*O*^-^) that is calculated from the data. Suppose a duplicate pair has *m* independent components, each of which is either ‘active’ (denoted by ‘1’) or ‘inactive’ (denoted by ‘0’). Let *U_11_* be the probability of a component being active in both genes; *U_01_* (or *U_10_*) is that of being inactive in gene *A* but active in gene *B* (or active in *A* but inactive in *B*); and *U_00_* be the probability of a component being inactive in both genes. Without loss of generality, one can assume *U_01_*=*U_10_*. Further, we claim *U_00_*=0 according to the *not-all-inactive constraint*, i.e., each component is functionally active at least in one duplicate copy. Together, *U_ij_*s can be concisely written as *U_11_*=1-2*U*, *U*_10_=*U*_01_=*U* and *U_00_* =0, respectively. In other words, with a probability of *2U*, a functional component is active in one duplicate but inactive in another one, and with a probability of 1-2*U*, a component is active in both duplicates.

Under the assumption that functional components of a gene are statistically independent and identical, one can derive *Q_t_*(*d_A_*, *d_B_*|*O*^-^) in term of the single component parameter (*U*) and the number (*m*) of functional components, that is,

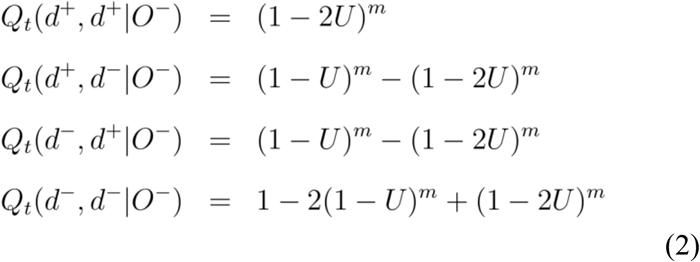

The rationale of Eq.(2) is as follows. Under the *m*-component model (*m*>1), two duplicate copies remain both dispensable only when each component is active in both duplicates (with a probability of 1-2*U*), which directly leads to the derivation of *Q_t_*(*d*^+^, *d*^+^|*O*^-^). Next we consider the (marginal) probability of dispensability (*d*^+^) conditional of the ancestral essentiality (*O*^-^), denoted by *Q_t_*(*d*^+^|*O*^-^). It appears that *Q_t_*(*d*^+^|*O*^-^)=(1-*U*)^*m*^ because the probability of a component for being active in one duplicate is given by (1-*U*). Since *Q_t_*(*d*^+^|*O*^-^)=*Q_t_*(*d*^+^, *d*^+^|*O*^-^)+*Q_t_*(*d*^+^, *d*^-^|*O*^-^), it is straightforward to obtain the second and third equations of Eq.(2). The last equation of Eq.(2) is derived by the sum of probabilities to be one.

### Rare neo-functionalization after duplication of dispensable gene

When a dispensable gene was duplicated, gene dispensability can be maintained through ancient genetic buffering and duplicate compensation. As a result, subfunctionalization becomes ineffective because the process of complementation between functional components (Force, et al. 1999) is difficult to achieve. While the neofunctionalization has been suggested for duplicate preservation (Chen, et al. 2010; Chen, et al. 2012), it is unlikely that both copes acquire new functions simultaneously. In this sense one can assume that

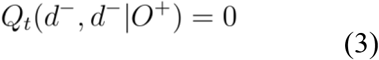

Intuitively speaking, this assumption holds well except for very ancient duplicates by independent new function acquiring that may be unrelated to the duplication event. Eq.(3) implies that any double-essential duplicate pairs (*d*^-^, *d*^-^) were virtually from the duplication of an essential gene (*O*^-^).

### Estimation of conditional probabilities

Based on Eqs.(2) and (3), we implement a computational procedure to calculate *Q_t_*(*d_A_, d_B_*|*O*^-^) and *Q_t_*(*d_A_, d_B_*|*O*^+^), respectively.

i. The proportion of dispensable single-copy genes is used as a proxy of *R_0_*.
ii. Applying *Q_t_*(*d*^-^, *d*^-^|*O*^+^)=0 to Eq.(1), leading to *P_t_*(*d*^-^,*d*^-^)=(1-*R_0_*) *Q_t_*(*d*^-^, *d*^-^|*O*^-^). Replacing *P_t_*(*d*^-^,*d*^-^) with its observed frequency *f_EE_*, we obtain

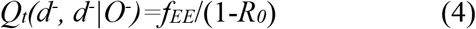
iii. The parameter *U* can be estimated by replacing the last equation of Eq.(2) of *Q_t_*(*d*^-^, *d*^-^|*O*^-^) by *f_EE_* (1-*R_0_* that is,

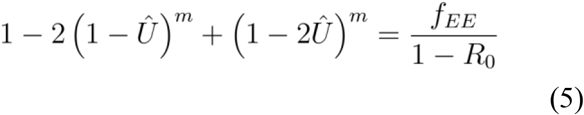

where *m,* the number of functional components, is treated as a known integer, i.e., *m*=2, 3,….
iv. When *U* is estimated by Eq.(5), the estimates of joint probabilities conditional of the ancestral essentiality (*O*^-^), *Q*(*d_A_,d_B_*|*O*^-^), can be obtained by replacing *U* with its estimate in Eq.(2). Eqs.(4) and (5) indicate that *f_EE_*, the frequency of double-essential duplicate pairs, determines the conditional probabilities of (*d_A_, d_B_*) after duplication of an essential gene (*O*^-^), provided that *R_0_* and *m* are treated as known constants.
v. It is straightforward to estimate *Q*(*d*^+^,*d*^+^|*O*^+^) by equating *P_t_*(*d*^+^, *d*^+^) with *f_DD_* in Eq.(1) in the case of *d_A_*=*d*^+^ and *d_B_*=*d*^+^, and replacing *Q*(*d*^+^,*d*^+^|*O*^-^) by its estimate. They are, respectively, given by

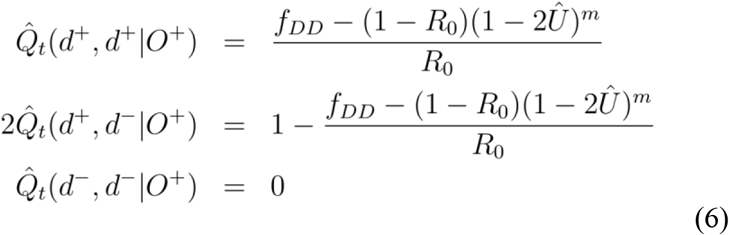

### Estimation of the sampling variances

The statistical property of those estimated conditional probabilities can be evaluated by calculating their approximate sampling variances. One approach invokes the bootstrapping algorithm. The other approach is the delta-method under a multinomial distribution of *f_DD_, f_DE_* and *f_EE_*, as briefly introduced below. Denote the function *g* by

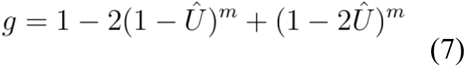

One can calculate the partial derivative of *g* by

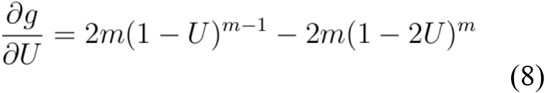

Hence, applying the delta-method in Eq.(5), we obtain

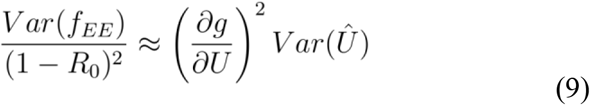

resulting in

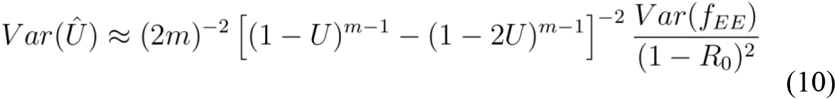

After some basic yet tedious algebras, the analytical forms for the sampling variances of those estimated conditional probabilities can be approximately derived in terms of the observed proportions *f_DD_, f_DE_* and *f_EE_*, as well as *N* (the number of duplicate pairs).

### Case-study: yeast duplicate pairs from whole genome duplications

We applied the new method to analyze 325 yeast duplicate pairs from the yeast WGD (whole-genome duplication) about 100 million years ago. The fitness dataset of single gene-deletions for yeast genes were from Gu et al. (2003). The fitness value *f_i_* is measured by the growth rate of the strain with gene *i* deleted relative to the average growth rate of wild strains under five growth media: YPD, YPDGE, YPE, YPG, and YPL. Similar to Gu et al. (2003), we classified these duplicate genes under study into four groups: The normal (*N*)-group is defined by *f_min_*>0.95, where *f_min_* is the smallest *f*-value for all five growth conditions, i.e., the deletion has a weak or no fitness effect in all conditions. The *M*-group is defined by 0.8<*f_min_*<0.95 for the moderate effect, and the *S*-group by 0.5<*f_min_*<0.8 for a strong effect. Finally, the *L*-group is defined by *f_min_*<0.5 that includes the lethal effect (*f*=0). Tentatively, a gene is classified as *d*^+^ if it belongs to the *N* or *M*-group, or *d*^-^ otherwise. Under this cutoff, the observed frequencies for patterns DD, DE and EE are *f_DD_*=0.803±0.022, *f_DE_*=0.179±0.021 and *f_EE_*=0.018±0.0074, respectively. In addition, the proportion of dispensable single-copy genes (0.605) is used as a proxy of *R_0_*.

We first examine how the number (*m*) of functional components may affect our analysis. Note that single functional component (*m*=1) is not allowed. Hughes and Liberles (2007) suggested that between *m*=2 and *m*=12 regulatory regions would be biologically realistic. Stark et al. (2017) suggested that it is unlikely that a gene would have in excess of *m*=20 functional components. Within the range of *m*=2-10, the effects of *m* on our analysis of yeast duplicate pairs are shown in **Fig. 1** (panel A or B). Overall it reveals little difference among cases of *m*=3 or more; only marginal differences appear when *m*=2, and all estimates are virtually the same between *m*=20 and *m*=∞. We therefore conclude that the effect of variable *m* is usually negligible.

**Fig. 1.**
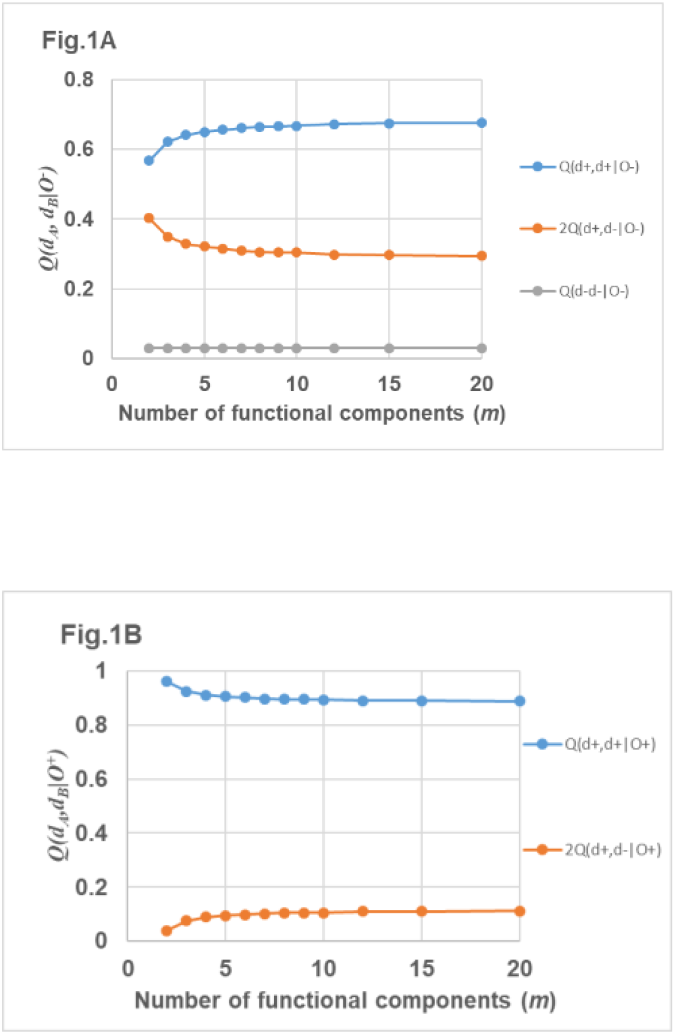
Estimated conditional probabilities of yeast WGD duplicate pairs plotting against the number of functional components *m*=2,…, 20. (A) *Q*(*d_A_, d_B_*|*O*^-^), predicted probabilities conditional of ancestral essentiality (*O*^-^). (B) *Q*(*d_A_, d_B_*|*O*^+^), predicted probabilities conditional of ancestral dispensability (*O*^+^).

The proportion of essential genes for the yeast duplicate pairs can be calculated by *P_E_*= *f_EE_*+*f_DE_*/2=0.103±0.012, which is significantly lower than the proportion of essential genes in single-copy genes (*P_E_*=0.395); p-value<10^-6^. Based on the newly-developed analysis, one can further estimate *P_E_* for genes duplicated from essential genes, which is given by *P_E_*(*O*^-^)=*Q_t_*(*d*^-^, *d*^-^|*O*^-^)+*Q_t_*(*d*^-^, *d*^-^|*O*^-^)=0.187±0.035 (*m*=6), whereas *P_E_* for genes duplicated from dispensable genes, which is given by *P_E_*(*O*^+^)= *Q_t_*(*d*^-^, *d*^-^|*O*^+^)+*Q_t_*(*d*^-^, *d*^-^|*O*^+^)=0.043±0.018 (*m*=6). Bootstrapping analysis shows that *P_E_*(*O*^-^) is significantly larger than by *P_E_*(*O*^+^); *p*-value<0.001. Therefore, a rapid increase of essentiality, or loss of dispensability may occur after the duplication of essential genes, approximately four-fold as fast as the process after the duplication of dispensable genes.

### Model assumptions

We acknowledge that the new probabilistic model for the evolution of genetic robustness after gene duplication is inevitably oversimplified. For instance, after duplication of an essential gene, the model assumes that two duplicate copies evolve under sub-functionalization, neglecting the chance of neo-functionalization or other mechanisms. Moreover, each functional component is assumed to undergo subfunctionalization independently. On the other hand, after duplication of a dispensable gene, the interactions between ancestral genetic buffering, duplicate compensation and neo-functionalization remain largely unknown. It should be noticed that some attributes of genetic mechanisms have not been taken into accounts, such as the effect of dosage balance, later-stage functional divergence, as described in Innan and Kondrashov (2010). Our future study will focus on the development of a more realistic model for the evolution of gene duplicates.

### Effects of the assumption *Q_t_*(*d*^-^, *d*^-^|*O*^+^)=0

A key assumption in our analyses is Eq.(3), that is, after duplication of a dispensable gene (*O*^+^), the chance for both duplicate copies to be essential is negligible. While it is biologically intuitive, it may cause some bias, especially for some very ancient duplicate pairs. We thus conducted a computer simulation study to examine this effect by letting *Q*(*d*^-^,*d*^+^|*O*^+^)=*q*, where *q* is a small positive value. Our preliminary simulation result shows that the estimation bias of those conditional probabilities is usually marginal, except for the case after an extremely long evolutionary span after gene duplication (not shown).

### Concluding remarks

This paper described a probabilistic model to explore the pattern of genetic robustness after the gene duplication, which makes a distinction between two evolutionary scenarios: duplication of an essential gene and duplication of a dispensable gene. This model was applied to yeast duplicate pairs from a whole-genome duplication, revealing that the proportion of essential genes for those duplicated from essential genes is approximately four-fold as high as that for those duplicated from dispensable genes. While more genome-wide single-gene deletion (knockout) data for duplicate pairs are available in different species, it is anticipated that the new model can provide some new insights about the role of duplicate compensation on genetic robustness.

